# *In Vitro* Prebiotic Potential of Purified Black Rice Oligosaccharides: Simulated Digestion Stability, Short-Chain Fatty Acid Production, and Probiotic Biofilm Formation

**DOI:** 10.64898/2026.02.10.705216

**Authors:** Shivangi Agrawal, Paramita Biswas, Susmita Mondal, Abinaya Balasubramanian, Sachin Maji, Sandip Shit, Satyabrata Ghosh, Satyahari Dey

## Abstract

Black rice oligosaccharides (BO) were extracted with 80% aqueous ethanol (v/v) and purified by charcoal–celite chromatography followed by dialysis using a 500 Da molecular weight cut-off (MWCO) membrane, yielding an oligosaccharide fraction with a degree of polymerisation (DP) between three and eight (DP3–DP8; 2.87 ± 0.29% w/w). MALDI-TOF MS showed sodium adduct ions from m/z 527 to 1330, and GC-MS analysis of hydrolysed samples identified glucose and galactose as the major monomers, while ketosyl residues were detected in the intact fraction by selective staining and are most plausibly attributed to fructosyl units based on cereal origin and DP distribution. BO showed high resistance to simulated salivary, gastric, and pancreatic digestion (only 2.08 ± 0.51%, 0.34 ± 0.03%, and 4.29 ± 0.73% hydrolysis, respectively) with approximately 93% remaining carbohydrate available for fermentation. All *Lactobacillus* strains showed positive prebiotic activity scores, with the highest response observed for *Lactobacillus rhamnosus* (1.165 ± 0.255) and *Lactobacillus plantarum* (0.980 ± 0.163). Fermentation produced metabolically relevant short-chain fatty acids (SCFA), mainly acetate (34.82 ± 2.08 mM), as well as strain-dependent propionate and butyrate levels. BO greatly promoted probiotic biofilm formation, with biomass reaching 391.33 ± 26.08% and viable cell counts of 9.01 ± 0.70 log CFU/mL relative to the control. Collectively, the results indicate that BO represents a digestion-resistant, hexose-based oligosaccharide series that is selectively utilised by probiotic lactobacilli, promotes SCFA production and enhances biofilm development. To our knowledge, this work is the first to combine structural profiling with *in vitro* functional evaluation of a purified, low-DP oligosaccharide fraction obtained from black rice.

**Highlights:**

- Purified oligosaccharides (DP3–DP8) were obtained from black rice using charcoal–celite chromatography followed by dialysis.
- Structural analysis confirmed that the oligosaccharides were hexose-based and composed mainly of glucose and galactose.
- Black rice oligosaccharides exhibited higher resistance to simulated gastric and intestinal digestion compared with starch.
- Positive prebiotic activity scores were observed due to selective utilisation by probiotic *Lactobacillus* strains.
- Fermentation of black rice oligosaccharides significantly increased short-chain fatty acid production.
- Purified oligosaccharides enhanced probiotic biofilm formation, indicating improved colonisation potential.

**Graphical abstract:** 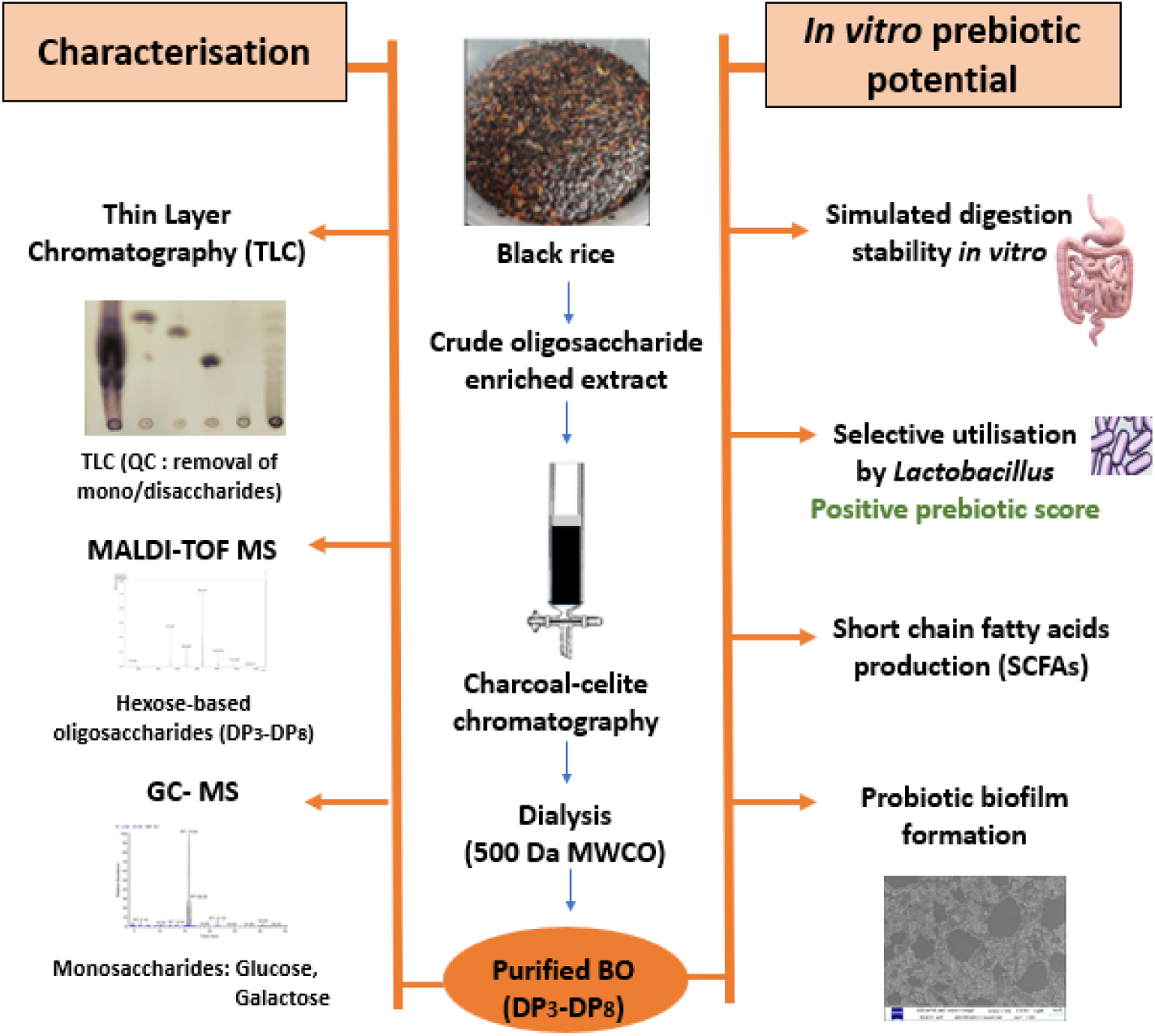

## Introduction

The rising incidence of metabolic diseases has generated considerable interest in functional dietary components that influence gut microbiota composition and activity (Li et al., 2021; Rashwan et al., 2024). Prebiotics are dietary components that escape digestion and are able to induce specific microbes in the gut, resulting in beneficial alterations in microbial activity and host physiology (Gibson and Roberfroid, 1995; Roberfroid, 2007). Non-digestible oligosaccharides are a key group of these carbohydrates because they largely escape digestion in the proximal gastrointestinal tract (GIT), thereby entering the colon unaltered and becoming available for preferential microbial fermentation. This fermentation can reshape microbial ecology and results in SCFA, principally acetate, propionate, and butyrate, which support colonocyte energy metabolism, epithelial barrier integrity, and immune regulation (Bindels et al., 2015; Playne & Crittenden, 2009; Morrison and Preston, 2016). Although fructooligosaccharides (FOS), inulin, and galactooligosaccharides (GOS) have been extensively characterised, there is increasing interest in cereal-derived oligosaccharides as sustainable and structurally diverse alternatives for functional food applications (Patel and Goyal, 2011).

Pigmented cereals such as black rice (*Oryza sativa* L.) contain complex carbohydrate matrices that include a low-molecular-weight soluble carbohydrate fraction (Abdel-Aal et al., 2006; Sompong et al., 2011). Most published work has examined whole-grain or bran-derived extracts rather than isolating and characterising the corresponding oligosaccharide components (Bae et al., 2017). Although small soluble oligosaccharides have been noted in black rice, their structural identity, degree of polymerisation, and monomeric composition have not been systematically defined. Moreover, while aqueous ethanol extraction is known to release soluble carbohydrates from cereal matrices, the composition and functional behaviour of a purified oligosaccharide fraction obtained after charcoal–celite chromatography and dialysis have not been established. To date, there is no report describing a structurally characterised, low-molecular-weight oligosaccharide fraction from black rice or evaluating its *in vitro* prebiotic functionality.

Resistance to hydrolysis by salivary, gastric, and pancreatic enzymes is a key requirement for prebiotic oligosaccharides (Roberfroid, 2007), and *in vitro* simulated digestion systems are commonly utilised to determine this property (Wichienchot et al., 2010). The digestibility under simulated gastrointestinal conditions is well known for reference non-digestible oligosaccharides, including FOS and inulin (Hernández-Hernández et al., 2019); however, analogous digestibility data for black rice oligosaccharides is lacking. After surviving gastrointestinal transit, oligosaccharides act as selective substrates for beneficial bacteria including *Lactobacillus* spp., whose fermentative metabolism yields acetate, propionate, and butyrate that support epithelial energy balance, barrier integrity, and immune regulation (Morrison and Preston, 2016). The SCFA acidification process creates an environment that enables probiotic bacteria to thrive while simultaneously preventing pathogenic bacteria from multiplying. While the fermentability and SCFA production potential of many cereal-derived oligosaccharides have undergone investigation (Harris et al., 2019; Tran et al., 2020), no functional investigations have delineated the fermentation profile of black rice oligosaccharides.

Prebiotics may also impact on probiotic biofilm formation, an ecological factor that increases mucosal adhesion, colonisation stability and pathogen exclusion (Lebeer et al., 2007; Jones and Versalovic, 2009). Despite its relevance to probiotic persistence in the gut environment, the effect of black rice-derived oligosaccharides on *Lactobacillus* spp. biofilm development remains poorly explored.

To address these knowledge gaps, the present study examined the extraction, purification, and structural profiling of black rice oligosaccharides using ethanol extraction, charcoal–celite chromatography, dialysis, thin-layer chromatography, MALDI-TOF MS, and GC-MS. Digestive resistance was assessed under simulated salivary, gastric, and intestinal conditions, while selective utilisation and SCFA production were quantified in *Lactobacillus* spp. cultures. The effect of the purified oligosaccharide fraction on probiotic biofilm formation was assessed quantitatively and microscopically. In this work, integrated approaches of structural characterisation, in vitro digestion, and functional assays were utilised for evaluation of the purified black rice oligosaccharides as digestion-resistant substrates promoting probiotic fermentation and biofilm formation, relevant for their role as cereal-derived oligosaccharide ingredients for gut health applications.

## 2. Materials and Methods

### 2.1 Plant Material

Black rice (*Oryza sativa* L., Manipuri cultivar) was procured from True Elements, Pune. The grains washed in distilled water and then dried in a hot air oven at 50 °C for 12–14 h before processing. The dried material was ground using a domestic mixer-grinder (Philips HL1643, India) and sieved to obtain flour with a particle size range of 150–500 µm. The resulting flour was stored in airtight glass containers at 4°C until further analysis.

### 2.2 Chemicals, Media, and Microorganisms

All reagents employed in the study were of analytical grade. Glucose, sucrose, raffinose, fructooligosaccharides (FOS), inulin, phenol, N-methyl-N-(trimethylsilyl)trifluoroacetamide (MSTFA), human salivary amylase, pancreatic amylase, and methoxyamine hydrochloride were procured from Sigma-Aldrich (USA). Potassium persulfate was purchased from HiMedia (India). Solvents such as ethanol, methanol, butanol, propanol and acetic acid as well as acids like nitric acid, sulfuric acid or hydrochloric acid were bought from Merck KGaA (Darmstadt, Germany). Indicators, common laboratory salts, silica plates, and Celite were also from Merck; activated charcoal was purchased from Loba Chemie (Mumbai, India). Dialysis tubing, MWCO 500 Da was purchased from Thermo Scientific (USA) and the 3,5-dinitrosalicylic acid (DNS) reagent was procured from SRL (India). The SCFA standards including acetic, butyric, and propionic acids were purchased from Dr. Ehrenstorfer GmbH (Germany).

All the components of culture media such as MRS broth and agar, peptone, tryptone, yeast extract, and crystal violet were purchased from HiMedia (India). The probiotic strains *Lactobacillus acidophilus* MTCC 10307, *Lactobacillus brevis* MTCC 1750, *Lactobacillus rhamnosus* MTCC 1408, *Lactobacillus plantarum* MTCC 2621, and *Lactobacillus casei* NCDC 357, and the enteric control strain *Escherichia coli* ATCC 15223 were used in all experiments.

### 2.3 Extraction and Purification of Black Rice Oligosaccharides (BO)

#### 2.3.1 Extraction

Black rice flour (100 g) was extracted with 80% (v/v) aqueous ethanol using a solvent-to-solute ratio of 5:1 (v/w) and maintained at 60 °C for 1 h under intermittent agitation. Following extraction, the suspension underwent centrifugation at 13,776 × g for 15 min in 50 mL Oak Ridge tubes, and the clarified supernatant was recovered. Ethanol was subsequently removed under reduced pressure using a rotary evaporator (Büchi R-210, Switzerland) operated at 55 °C, 150 mbar, and 50 rpm. The remaining aqueous phase was freeze-dried (Eyela FDU-1200, Japan) to obtain a crude extract enriched in oligosaccharides.

#### 2.3.2 Charcoal–Celite Chromatography and Dialysis

The crude extract was dissolved in distilled water and loaded onto an activated charcoal–celite column (1:1, w/w; 30 cm × 4 cm i.d.). The sample-to-adsorbent ratio was maintained at 1:5 (w/w) following Mondal et al. (2022). A stepwise elution procedure was carried out with water followed by increasing concentrations of aqueous ethanol (5, 10, 15, 50, and 90% v/v). Fractions were concentrated to remove ethanol and freeze-dried Fractions enriched in low-molecular-weight oligosaccharides, as confirmed by TLC and sugar profiling, were combined and subjected to dialysis against distilled water at 4 °C for 48 h using a 500 Da MWCO membrane. The dialysed retentate was freeze-dried to yield purified BO.

### 2.4 Carbohydrate Estimation and TLC Profiling

#### 2.4.1 Total Carbohydrate Content

Total carbohydrate content was measured by a microplate-adapted phenol–sulfuric acid method (Dubois et al., 1956). Aliquots of the samples (20 µL) were combined with 20 µL of 5% (w/v) phenol reagent and 200 µL of concentrated H₂SO₄. After a reaction time of 10 min, the absorbance was measured at 490 nm. Glucose was employed for the preparation of the calibration curve.

#### 2.4.2 Reducing Sugar Content

Reducing sugar levels were assessed utilizing the DNS assay (Miller, 1959). The sample and DNS reagent were mixed in equal volumes and were placed in a boiling water bath for 10 min, followed by the addition of potassium sodium tartrate at a 1:3 (v/v) ratio. Absorbance was measured at 540 nm. Glucose was employed as the reference standard.

#### 2.4.3 TLC Profiling of Soluble Carbohydrates

Samples were spotted approximately 1 cm from the lower edge of silica gel 60 F₂₅₄ TLC plates and resolved using a solvent system consisting of acetic acid: n-butanol: water (1:2:1, v/v/v). After solvent evaporation, the plates were subjected to treatment with orcinol–sulfuric acid reagent and thermally developed at 80 °C to enable visualisation of carbohydrate bands.

### 2.5 Structural Characterisation

#### 2.5.1 Molecular Weight Profiling by MALDI-TOF MS

Lyophilised BO (1 mg) was dissolved in 0.01 M NaCl and mixed with a 2,5-dihydroxybenzoic acid (DHB) matrix solution (5 mg in 50% (v/v) ethanol) in a 1:1 (v/v) ratio. An aliquot (2 µL) of the prepared mixture was deposited onto a stainless-steel target plate, allowed to air-dry, and subjected to analysis using a Bruker Ultraflextreme MALDI-TOF/TOF instrument (Bruker Daltonik GmbH, Germany) operated in reflectron positive-ion mode over an m/z range of 100–2000. Detected oligosaccharide signals were assigned on the basis of sodium adduct formation ([M + Na]⁺).

#### 2.5.2 Monosaccharide Composition Analysis

Purified BO (2–3 mg) was subjected to acid hydrolysis using 4 M TFA at 120 °C for a duration of 2–3 h. Residual TFA was eliminated through repeated co-evaporation with methanol. The dried hydrolysates were derivatised for GC–MS analysis using 3% (w/v) methoxyamine hydrochloride in pyridine at 37 °C for 2 h, followed by the addition of 70 µL MSTFA and incubation of the reaction mixture for 30–40 min.

GC–MS analysis was conducted using a Thermo system fitted with a TR-5MS column (30 m × 0.25 mm, 0.25 µm). The oven temperature program escalated from 50 °C to 130 °C at a rate of 3 °C/min, followed by a 2 min hold. The inlet temperature was maintained at 230 °C, and spectra were recorded from m/z 50–400 at a split ratio of 1:20. Monosaccharide identities were confirmed through comparison with derivatised reference standards and database entries from the NIST mass spectral library.

The presence of ketose residues was confirmed by the urea–phosphoric acid test (Wise et al., 1955), wherein TLC plates were sprayed with the reagent and placed at 110 °C for 5–7 min to allow development of a characteristic colour.

### 2.6 *In Vitro* Non-Digestibility Assay

BO solutions (1% w/v) were incubated separately with salivary α-amylase (10 U mL⁻¹) or pancreatic α-amylase (10 U mL⁻¹) in 20 mM sodium phosphate buffer (pH 7.0). To simulate gastric digestion, BO solutions were exposed to 0.01 N HCl (pH 2.0). All digestion assays were conducted at 37 °C over a period of 6 h, with samples withdrawn at 0 h and 6 h. Enzymatic activity was halted by thermal inactivation at 100 °C for 5 min. Acid-treated samples were subsequently neutralised to pH 7.0 using NaOH prior to further analysis. Starch (digestible control) and FOS (prebiotic reference) were processed in parallel under identical conditions. Total carbohydrate and reducing sugar contents were determined by the phenol–sulfuric acid and DNS assays, respectively. The extent of hydrolysis was calculated following Wichienchot et al. (2010), as shown below using Eq. (1): Percentage hydrolysis (%) = (Reducing sugar released × 100) / (Total sugar content − Initial reducing sugar content). (1) Where the reducing sugar released represents the difference between the final and initial reducing sugar concentrations.

### 2.7 Prebiotic Activity Assay

Probiotic strains (*L. plantarum, L. acidophilus, L. brevis, L. rhamnosus,* and *L. casei*) were separately inoculated (1% v/v) into carbohydrate-free basal MRS medium supplemented with BO (1% w/v), glucose (1% w/v; growth control), or FOS (1% w/v; reference prebiotic) as the sole carbon source. Cultures were maintained at 37 °C, and viable counts were determined on MRS agar at 0 h and 24 h.

*Escherichia coli* was grown in M9 medium and inoculated (1% v/v) into M9 minimal medium supplemented with BO (1% w/v), glucose (1% w/v; growth control), or FOS (1% w/v; reference prebiotic). Enumeration was performed on nutrient agar at 0 h and 24 h.

The prebiotic activity score was calculated according to Huebner et al. (2007), using Eq. (2): Prebiotic activity score = {(probiotic log CFU/mL on the prebiotic at 24 h − probiotic log CFU/mL on the prebiotic at 0 h) / (probiotic log CFU/mL on glucose at 24 h − probiotic log CFU/mL on glucose at 0 h)} − {(enteric log CFU/mL on the prebiotic at 24 h − enteric log CFU/mL on the prebiotic at 0 h) / (enteric log CFU/mL on glucose at 24 h − enteric log CFU/mL on glucose at 0 h)} (2)

### 2.8 SCFA Quantification by HPLC

Cell-free fermentation broths for SCFA analyses were collected after 48 h of growth in BO-supplemented basal MRS or M9 medium (for *Lactobacillus* strains and *E. coli*, respectively). The cultures were clarified by centrifugation (10,000 × g for 10 min) and the supernatant was filtered through sterile 0.22 µm membrane filters.

The SCFA (acetic, propionic, and butyric acids) were analysed on an Agilent 1200 HPLC system coupled with a Hypersyl Gold C18 column (250 mm × 4.6 mm, 5 µm) maintained at 30 °C, following the protocol of Mitra et al. (2016). Mobile phase A was composed of NaH₂PO₄ (20 mM) supplemented with phosphoric acid to adjust the pH to 2.2, and mobile phase B contained acetonitrile. Chromatographic separations were conducted at a flow rate of 1.25 mL min⁻¹, with 20 µL sample injections and UV detection at 210 nm using a diode array detector, employing gradient elution. A mixed SCFA standard was analysed under identical conditions, and individual calibration curves were used for quantification.

### 2.9 Quantitative and Qualitative Probiotic Biofilm Analysis

The influence of BO on probiotic biofilm formation was determined in a mixed *Lactobacillus* culture as per O’Toole (2011), with slight modifications. BO (1% w/v) and FOS (1% w/v) were prepared in carbohydrate-free basal MRS medium and sterilised by filtration through 0.22 µm membrane filters. Carbohydrate-free basal MRS medium without any added carbohydrate served as the negative control, whereas FOS (1% w/v) served as the reference prebiotic control. A mixed culture of *L. acidophilus*, *L. rhamnosus*, *L. brevis*, *L. plantarum*, and *L. casei* (1:1:1:1:1) was adjusted to approximately 10⁸ CFU/mL and dispensed into 96-well plates (200 µL per well) or onto sterile glass coverslips in 24-well plates. Plates were incubated statically at 37 °C for 48 h, and all experiments were conducted in biological triplicate.

#### 2.9.1 Crystal Violet Biofilm Quantification

After incubation, non-adherent cells were aspirated, and the wells were carefully washed with PBS (pH 7.2) to eliminate loosely attached bacteria. Biofilms were stabilised by fixation with methanol for 20 min, followed by staining with 0.1% (w/v) crystal violet for 15 min and subsequent washing with deionised water. Biofilm morphology was examined using a bright-field microscope (Leica DMi8, Germany). The retained crystal violet was released by treatment with 33% (v/v) glacial acetic acid (100 µL per well), and absorbance values were recorded at 590 nm. Biofilm formation was calculated relative to the carbohydrate-free control using Eq. (3):

Biofilm formation (%) = (Absorbance₅₉₀, treatment / Absorbance₅₉₀, control) × 100 (3)

#### 2.9.2 Enumeration of Viable Bacteria in Biofilms

Unstained biofilms were resuspended in PBS, vortexed to dislodge adherent bacterial cells and serially diluted. Ten microlitre aliquots were plated on MRS agar. Plates were incubated at 37 °C for 24 h, and viable counts were expressed as log CFU/mL of the biofilm solution.

#### 2.9.3 Scanning Electron Microscopy (SEM)

Biofilms formed on glass coverslips were gently rinsed with PBS and chemically fixed using 4% (v/v) glutaraldehyde for 2 h at ambient temperature. The fixed samples were washed twice with PBS and subjected to graded dehydration through 35%, 50%, 75%, and 95% ethanol for 10 min at each step, followed by three successive treatments with 100% ethanol for 15 min each. The coverslips were subsequently vacuum-dried, mounted onto aluminium stubs using conductive carbon tape, and coated with a thin layer of gold by sputtering. Biofilm surface architecture was visualised using a scanning electron microscope (Zeiss, Germany).

### 2.10 Statistical Analysis

All assays were conducted in three independent biological replicates, and data are presented as mean ± SEM. Statistical analyses were performed using GraphPad Prism version 5.0 (GraphPad Software Inc., USA). Group differences were tested by one-way ANOVA, with Dunnett’s multiple comparisons test (versus control) applied for multiple comparisons within each strain. Statistical significance was set at p < 0.05.

## 3. Results and Discussion

### 3.1. Extraction And Purification of Black Rice Oligosaccharides (BO)

Aqueous ethanol extraction is widely used for isolating low-molecular-weight oligosaccharides from cereal matrices because it solubilises small carbohydrates while limiting the co-extraction of starch, proteins, and other macromolecules (Ekvall et al., 2007). Extraction of black rice flour with 80% (v/v) aqueous ethanol yielded a crude oligosaccharide-rich fraction of 4.95 ± 0.33% (w/w). The TLC profile showed distinct bands corresponding to mono-, di- and low-degree-of-polymerisation (DP) oligosaccharides with no polysaccharide streaking. This confirmed that 80% (v/v) aqueous ethanol effectively restricted starch dissolution. These observations align with earlier reports indicating that increasing ethanol concentration reduces solvent polarity and suppresses endogenous hydrolases such as α-galactosidase and invertase (Balto et al., 2016; Johansen et al., 1996).

Purification through charcoal–celite chromatography produced a characteristic ethanol-dependent desorption pattern. Monosaccharides were primarily recovered in water and 5% (v/v) aqueous ethanol, disaccharides in 10% (v/v) aqueous ethanol, and DP3–DP8 oligosaccharides were most enriched in the 15% and 50% (v/v) aqueous ethanol fractions (Fig. 1B and Fig. 1C). This gradual release of carbohydrates was in accordance with a reported reduction of sugar–charcoal interactions at increased ethanol concentration (Whistler and Durso, 1950; Kuhn and Filho, 2010). Analogous elution patterns are also found for raffinose-family oligosaccharides and cereal fructans (Wienberg et al., 2022).

**Fig. 1.**
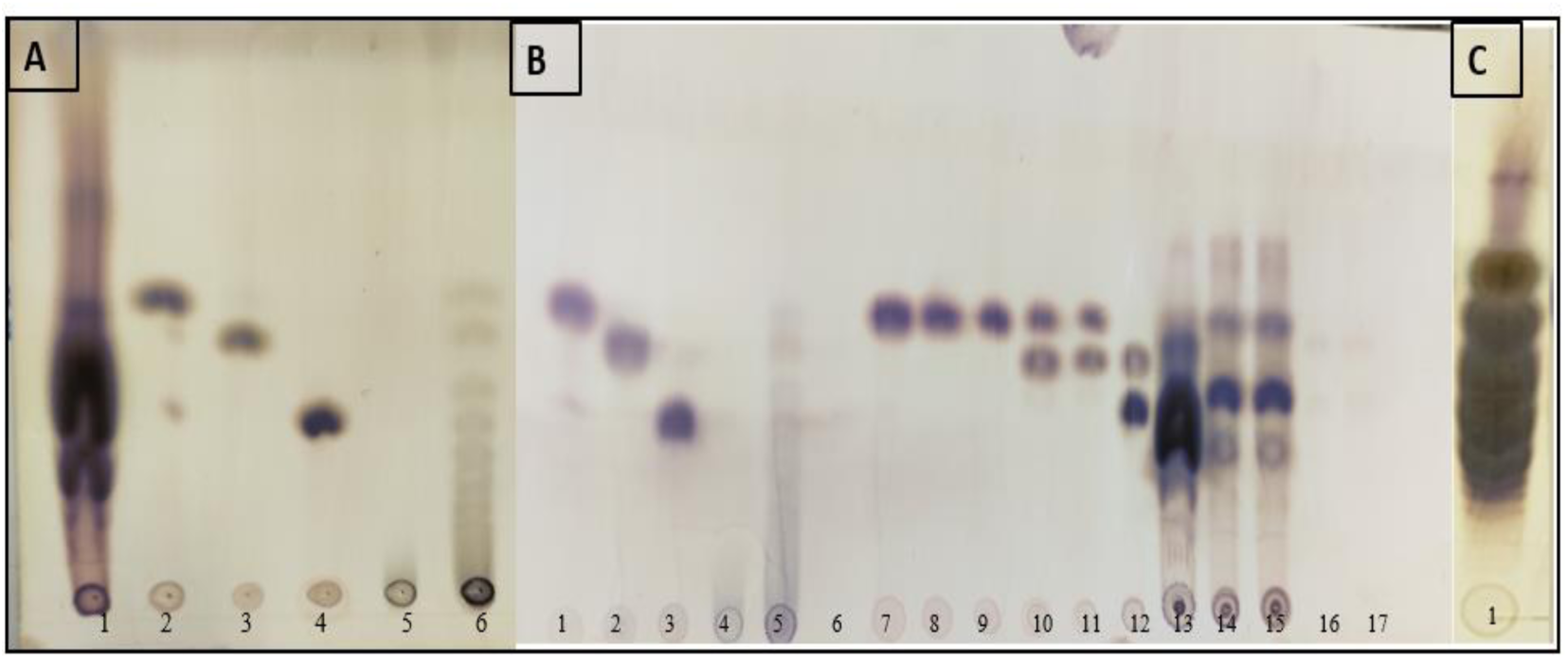
Thin-layer chromatographic analysis of the extraction and purification steps for black rice oligosaccharides. (A)Total sugars extracted: Crude carbohydrate extract obtained using 80% (v/v) aqueous ethanol (lane 1), while reference components were loaded as follows: glucose (lane 2), sucrose (lane 3), raffinose (lane 4), inulin (lane 5), and FOS (lane 6). (B) TLC of fractions obtained during charcoal–celite purification. Lanes 1–5 contain reference components: glucose (lane 1), sucrose (lane 2), raffinose (lane 3), inulin (lane 4), and FOS (lane 5). Lane 6 represents the initial column flowthrough, while lane 7 represents the water wash. Duplicate eluates collected with 5%, 10%, 15%, 50%, and 90% (v/v) aqueous ethanol are shown in lanes 8–9, 10–11, 12–13, 14–15, and 16–17, respectively. (C) Lane 1: dialysed oligosaccharide fraction obtained by pooling the 15% and 50% (v/v) aqueous ethanol eluates.

The recovery summary is presented in Table 1. From the crude extract, the charcoal–celite step yielded 1.06 ± 0.48% (5% ethanol eluent), 0.61 ± 0.23% (10% ethanol eluent), 1.08 ± 0.39% (15% ethanol eluent), and 2.12 ± 0.37% (50% ethanol eluent). The 90% (v/v) aqueous ethanol fraction did not contain detectable carbohydrate.

**Table 1.** Recoveries of oligosaccharides from 100 g black rice flour at each purification stage.

| Purification stage | Pre-dialysis | Charcoal–celite chromatography (stepwise aqueous ethanol elution) |  |  |  |  | Post-dialysis |
| --- | --- | --- | --- | --- | --- | --- | --- |
|  |  | 5 % | 10 % | 15 % | 50 % | 90 % |  |
| Dry weight (%) | 4.95 ± 0.33 | 1.06 ± 0.48 | 0.61 ± 0.23 | 1.08 ± 0.39 | 2.12 ± 0.37 | — | 2.87 ± 0.29 |

Residual mono- and disaccharides were removed by dialysis against distilled water using a 500 Da MWCO membrane, yielding a purified BO fraction of 2.87 ± 0.29%. Post-dialysis TLC confirmed the complete loss of monosaccharide and disaccharide bands. The extraction efficiency and purification behaviour observed in the present study are consistent with reports on pearl millet (Mondal et al., 2022), lupin (Martínez-Villaluenga et al., 2005), soybean (Hu et al., 2017), and pea cultivars (Jones et al., 1999).

In conclusion, the extraction and purification strategy combining 80% (v/v) aqueous ethanol, charcoal–celite chromatography and 500 Da MWCO dialysis yielded a structurally clean BO fraction free of mono- and disaccharides as well as polysaccharides and enriched in DP3–DP8 oligosaccharides. Although activated charcoal does not provide fine resolution of individual oligosaccharide species due to its heterogeneous microporosity, the method was robust and reproducible and yielded sufficient material for subsequent structural (MALDI-TOF MS and GC–MS) and functional assessments.

### 3.2 Structural Characterisation: Profiling and Monosaccharide Composition

Analysis by MALDI-TOF MS of the purified BO fraction revealed a series of sodium adducts from m/z 527 to approximately m/z 1330, corresponding to oligosaccharides with a degree of polymerisation between 3 and 8 (Fig. 2A). The DP range of 3–8 observed for BO fits within the window known to be associated with non-digestible carbohydrate fractions with prebiotic potential and selective microbial utilisation (Bindels et al., 2015). The mass of each successive ion increased by ∼162 Da, the size of a single hexose unit. This progressive sequence indicates the existence of a homologous series of hexose oligosaccharides. Although MALDI-TOF does not report linkage positions, this continuous 162 Da increment indicates a regular hexose-elongation pattern consistent with plant oligosaccharide biosynthesis. Similar DP distributions have been reported for prebiotic galactosyl saccharides in onion, garlic, mung bean, soybean, oat and other cereal matrices (Ritsema and Smeekens, 2003; Wichienchot et al., 2010), suggesting that BO contains nutritionally relevant oligosaccharide species with structural complexity typical of recognised prebiotic carbohydrates.

**Fig. 2.**
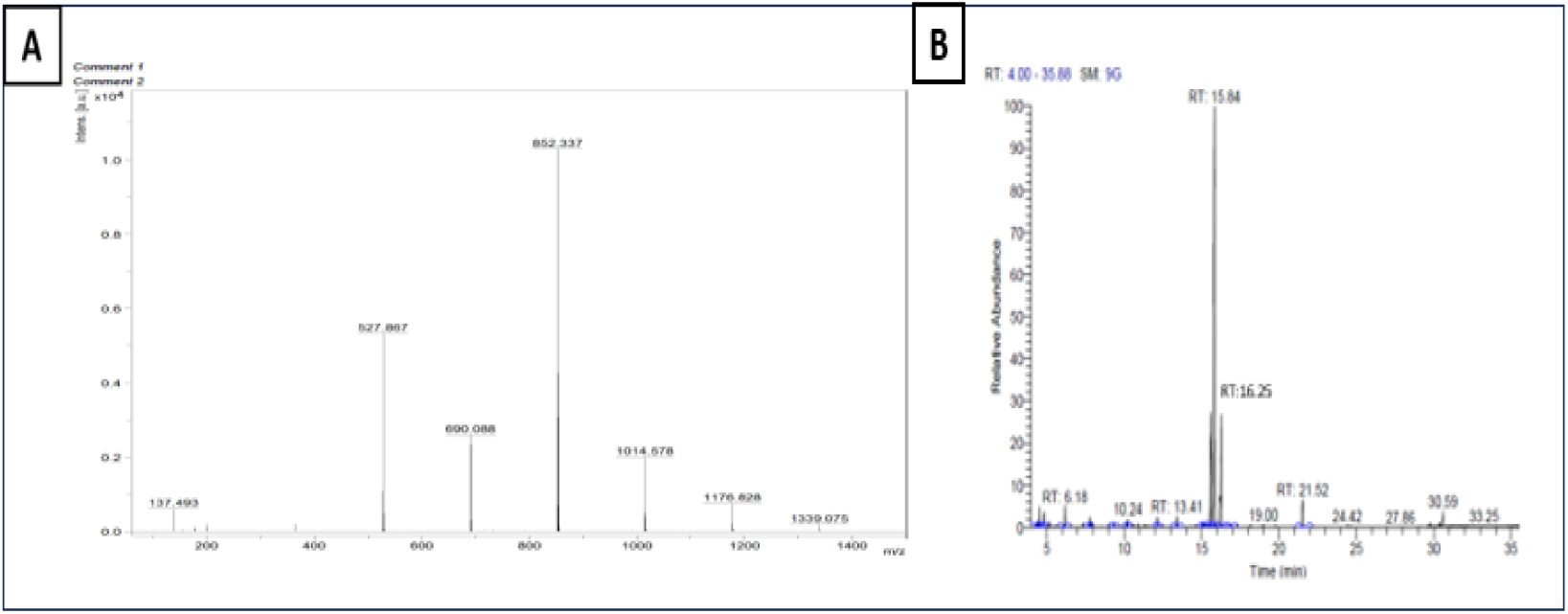
(A) MALDI-TOF mass spectrum of the dialysed oligosaccharide fraction isolated from black rice by pooling the 15% and 50% (v/v) aqueous ethanol eluates following charcoal–celite chromatography. (B) GC-MS chromatogram of trimethylsilylated monosaccharides released after acid hydrolysis of purified BO.

Monosaccharide composition analysis of acid-hydrolysed BO identified glucose and galactose as the predominant residues. GC–MS of the derivatised hydrolysate showed two major peaks at RT 15.84 and 16.25 min, with fragmentation patterns consistent with glucose and galactose, respectively (Fig. 2B). This compositional profile aligns with earlier reports indicating that cereal-derived oligosaccharides commonly contain glucosyl and galactosyl units as principal monomeric components (Henry and Saini, 1989). The ketose-specific assay produced a positive response for the intact BO fraction, suggesting the presence of ketose-reactive residues within the oligosaccharide fraction. Although this assay does not identify individual ketose sugars, the DP3–DP8 distribution observed by MALDI-TOF MS and the characteristic 162 Da mass increments support a hexose-based oligosaccharide series, but do not on their own confirm fructosyl-containing structures. The lack of fructose in the hydrolysed, derivatised samples is therefore not necessarily surprising as under harsh conditions of hydrolysis and derivatisation, it is possible that recovery of fructofuranosyl units may be limited by dehydration or fragmentation (Whistler and BeMiller, 1961; Aspinall, 1983). Such an observation can perhaps be related to some cereal systems in which fructofuranosyl-oligosaccharides have been reported (Henry and Saini, 1989; Peterbauer and Richter, 2001), but further structural confirmation would be needed to gain definitive support.

### 3.3 *In Vitro* Non-Digestibility Assay

Prebiotic classification requires resistance to degradation by enzymes and acids in the upper GIT. The digestive stability of BO was assessed using simulated salivary, gastric, and intestinal digestion models and compared to starch as the digestible control and FOS serving as the prebiotic reference.

During simulated salivary digestion, BO demonstrated 2.08 ± 0.51% hydrolysis, which was comparable to FOS (2.79 ± 0.44%) and substantially lower than starch (31.6 ± 2.07%), as observed in the present study (Fig. 3A). These results are consistent with previous work on pitaya, *Ganoderma* and dragon fruit oligosaccharides which generally exhibited less than 7% degradation by salivary α-amylase (Wichienchot et al., 2010). Taken together, these findings indicate that BO is not orally degradable and can transit to the stomach substantially unchanged.

**Fig. 3.**
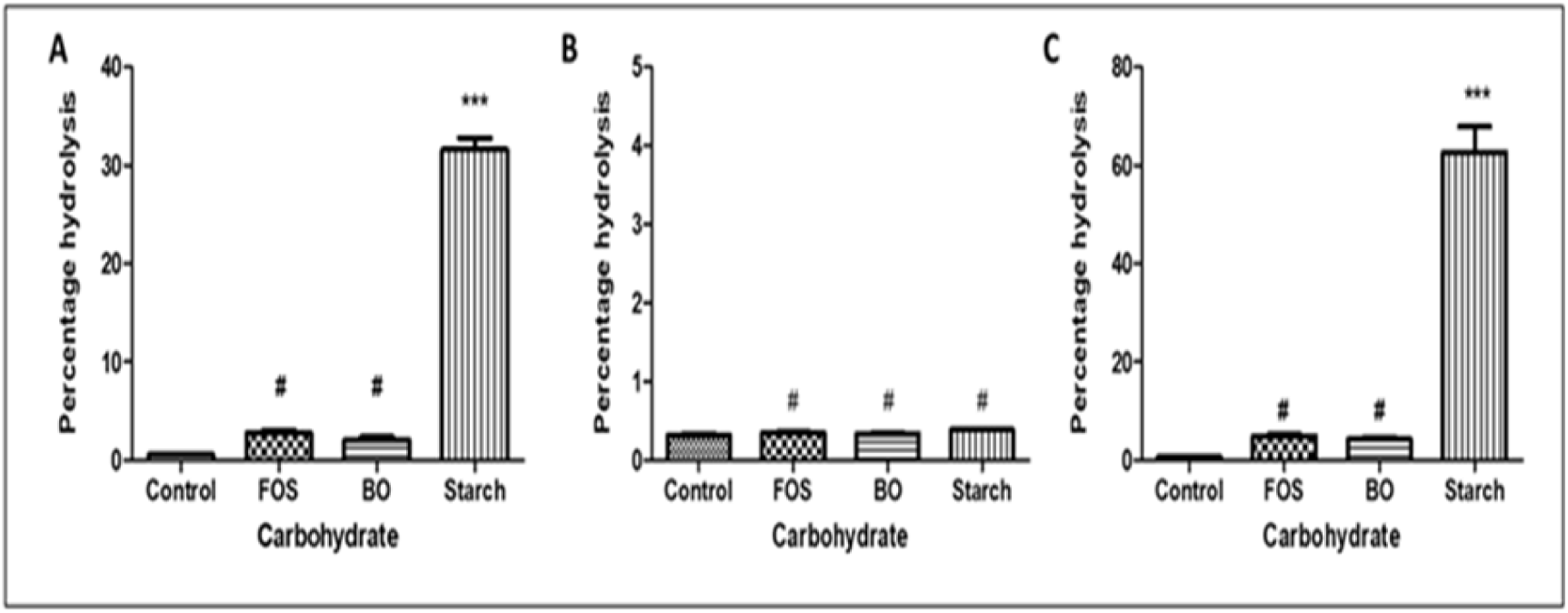
In vitro digestive stability of black rice oligosaccharides (BO) compared with fructooligosaccharides (FOS) and starch under simulated gastrointestinal conditions. Hydrolysis profiles following exposure to (A) salivary α-amylase, (B) simulated gastric conditions, and (C) pancreatic α-amylase are shown. Results are expressed as mean ± SEM from three independent experiments (n = 3). Differences among groups were analysed by one-way ANOVA followed by Dunnett’s multiple comparisons test (versus control). *p < 0.05, **p < 0.01, ***p < 0.001; # indicates not significant (p > 0.05) versus the control.

Hydrolysis under simulated gastric conditions was also low irrespective of the substrates. BO had 0.34 ± 0.03% hydrolysis, similar to FOS (0.36 ± 0.02%) and starch (0.39 ± 0.02%) (Fig. 3B). This high acid tolerance is in agreement with the literature. Pitaya oligosaccharides remain more than 96% intact under these conditions as well (Wichienchot et al., 2010). These data imply that BO may lack acid-labile glycosidic linkages and remains stable during gastric transit.

In simulated intestinal digestion with pancreatic α-amylase, BO showed high resistance (4.29 ± 0.73% of hydrolysis) comparable to FOS (4.98 ± 0.73%) and significantly lower than starch (62.67 ± 9.07%) (Fig. 3C). Similar resistance has been found with dragon fruit oligosaccharides and other cereal based non-digestible carbohydrates, which pass through pancreatic digestion more than 90% undigested (Bae et al., 2017; Pirkola et al., 2023). The relatively low digestibility of BO indicates an absence of easily hydrolysable α-(1→4) and/or α-(1→6) linkages which is also consistent with glycosidic configurations that are recognized to provide resistance towards human amylases, although conclusive linkage determinations necessitate additional structural analysis.

Across all digestive phases, approximately 93% of BO remained unhydrolysed. This resistance is comparable to that reported for well-established prebiotics, including inulin, FOS, and plant-derived GOS, which typically remain 88–100% intact during passage through the proximal gastrointestinal region (Cummings et al., 2001). Collectively, these findings indicate that BO can pass through the upper GI tract essentially intact, thereby fulfilling a key physiological criterion for prebiotic functionality.

### 3.4 Prebiotic Activity Assay

The selective utilisation of BO by probiotic bacteria was evaluated using the prebiotic activity score, which integrates probiotic growth promotion with suppression of enteric bacteria. All tested *Lactobacillus* strains exhibited positive prebiotic activity scores, indicating preferential utilisation of BO over the enteric control organism (*E. coli*). Among the strains examined, *L. rhamnosus* recorded the highest score (1.165 ± 0.255), followed by *L. plantarum* (0.980 ± 0.163) and *L. casei* (0.944 ± 0.076) (Fig. 4). *L. acidophilus* (0.770 ± 0.077) and *L. brevis* (0.469 ± 0.088) also demonstrated measurable utilisation of BO, although to a lesser extent.

**Fig. 4.**
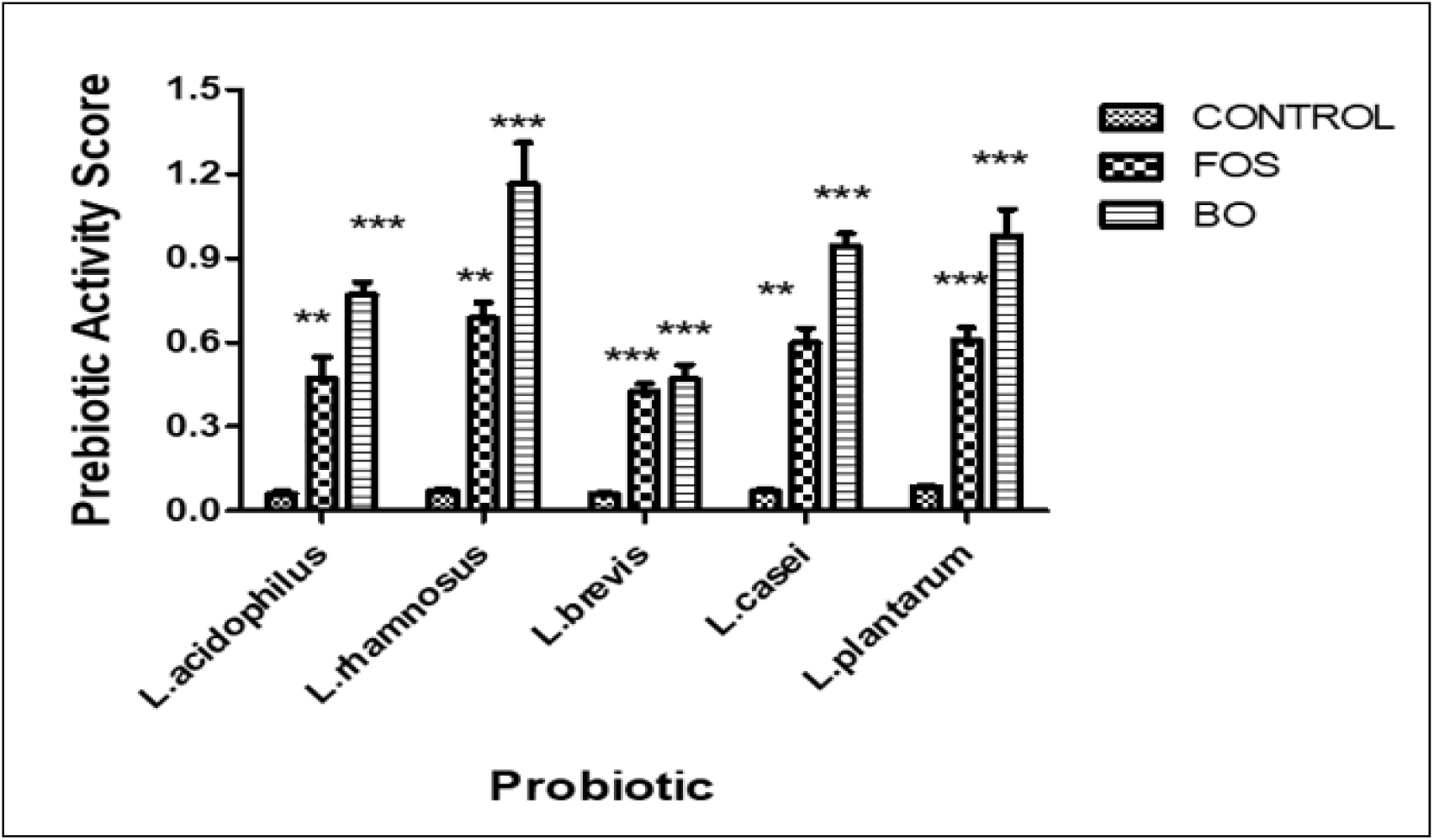
Prebiotic activity scores of black rice oligosaccharides (BO) across different *Lactobacillus* strains. Data are reported as mean ± SEM (n = 3). Differences among treatments within each strain were analysed by one-way ANOVA followed by Dunnett’s multiple comparisons test (versus control). *p* < 0.05, p < 0.01, *p* < 0.001; # indicates not significant (*p* > 0.05) versus the control.

The observed utilisation hierarchy (*L. rhamnosus* > *L. plantarum* > *L. casei* > *L. acidophilus* > *L. brevis*) reflects strain-specific differences in carbohydrate transport systems and intracellular glycosidase repertoires, which govern the metabolism of β-linked and structurally complex oligosaccharides (Andersen et al., 2012; Gänzle & Follador, 2012).

The prebiotic activity scores obtained for BO were comparable to, and in some cases higher than, those reported for established prebiotics such as FOS and GOS (typically ranging from ∼0.5–1.1, depending on strain and assay conditions) (Huebner et al., 2007). The higher scores observed for *L. casei* and *L. rhamnosus* indicate efficient and selective utilisation of BO, consistent with performance reported for conventional prebiotic substrates. Alongside its resistance to upper gastrointestinal digestion, these findings substantiate BO as a digestion-resistant carbohydrate with selective fermentability by lactobacilli, aligning with essential functional characteristics anticipated of prebiotics.

### 3.5 SCFA Quantification by HPLC

The SCFA generated from BO fermentation were determined in five *Lactobacillus* strains after 48 h incubation (Table 2). Acetic acid was the most abundant metabolite in all strains, and its maximum concentration was attained by *L. acidophilus* (34.82 ± 2.08 mM), followed by *L. plantarum* (30.80 ± 1.40 mM) and *L. rhamnosus* (28.23 ± 1.70 mM). *L. plantarum* exhibited the highest concentration of propionate (12.40 ± 0.64 mM), while the highest butyrate concentration was produced by *L. brevis* (5.31 ± 0.06 mM). The total SCFA output followed the order: *L. acidophilus* (47.91 ± 2.84 mM) > *L. plantarum* (46.30 ± 3.84 mM) > *L. rhamnosus* (43.10 ± 2.08 mM) > *L. casei* (40.57 ± 3.40 mM) > *L. brevis* (31.11 ± 2.60 mM).

**Table 2.** Production of SCFAs by *Lactobacillus* strains during fermentation of BO.

| SCFA (mM) | *L. acidophilus* | *L. brevis* | *L. casei* | *L. plantarum* | *L. rhamnosus* |
| --- | --- | --- | --- | --- | --- |
| Acetic acid | 34.82 ± 2.08 | 19.39 ± 0.74 | 26.25 ± 0.89 | 30.8 ± 1.4 | 28.23 ± 1.7 |
| Propionic acid | 8.9 ± 0.58 | 6.21 ± 1.8 | 10.24 ± 2.3 | 12.4 ± 0.64 | 11.93 ± 0.26 |
| Butyric acid | 4.19 ± 0.18 | 5.31 ± 0.06 | 4.08 ± 0.21 | 3.1 ± 1.8 | 2.94 ± 0.12 |

Values denote averages from three independent experiments, with SEM indicating
variability.

These strain-specific SCFA profiles indicate that, although BO was metabolised by all tested strains, carbon flux through acetate-, propionate-, and butyrate-producing routes differed markedly among strains. Similar substrate-dependent variation in SCFA output has been reported for lactobacilli grown on galactooligosaccharides, laminarin-derived oligosaccharides, and curdlan, and is generally attributed to differences in carbohydrate-active enzyme repertoires and central carbon metabolism (Chamberlain et al., 2022; Dong et al., 2023).

A disparity between prebiotic activity scores and SCFA production was also observed. *L. rhamnosus* exhibited the highest prebiotic activity score but only moderate SCFA production, indicative of a preference to direct BO-derived carbon toward biomass formation rather than metabolite secretion. *L. acidophilus* showed more limited growth responses but the highest total SCFA production, largely due to acetate accumulation. Similar decoupling between microbial growth and metabolite production has been reported previously and is attributed to strain-dependent carbon partitioning and regulation of fermentative pathways (Hamaker and Tuncil (2014)

The physiological significance of SCFA is well documented. Acetate plays a role in anti-inflammatory signalling and appetite regulation, propionate is associated with reduced hepatic lipogenesis and improved lipid metabolism, and butyrate serves as the primary energy source for colonocytes while promoting intestinal epithelial barrier function (Den Besten et al., 2013; Hamer et al., 2008). Notably, the simultaneous production of acetate, propionate, and butyrate from BO is of interest, as cereal-derived oligosaccharides frequently yield predominantly acetate-biased fermentation profiles. Thus, the SCFA profile observed in this study confirms the potential of BO as a metabolically favourable prebiotic substrate capable of modulating colonic fermentation beyond acetate-dominant outcomes.

### 3.6 Quantitative and Qualitative Probiotic Biofilm Analysis

The ability of BO to promote biofilm formation by *Lactobacillus* spp. was evaluated using bright-field microscopy, scanning electron microscopy (SEM), and quantitative assays. Bright-field micrographs (Fig. 5A1–C1) showed that BO supported greater surface-associated growth than FOS or the control. BO-treated cultures displayed dense cell clusters and a more continuous biofilm coverage, whereas FOS-treated cultures formed moderately developed biofilms and the control exhibited sparse, discontinuous attachment.

**Fig. 5.**
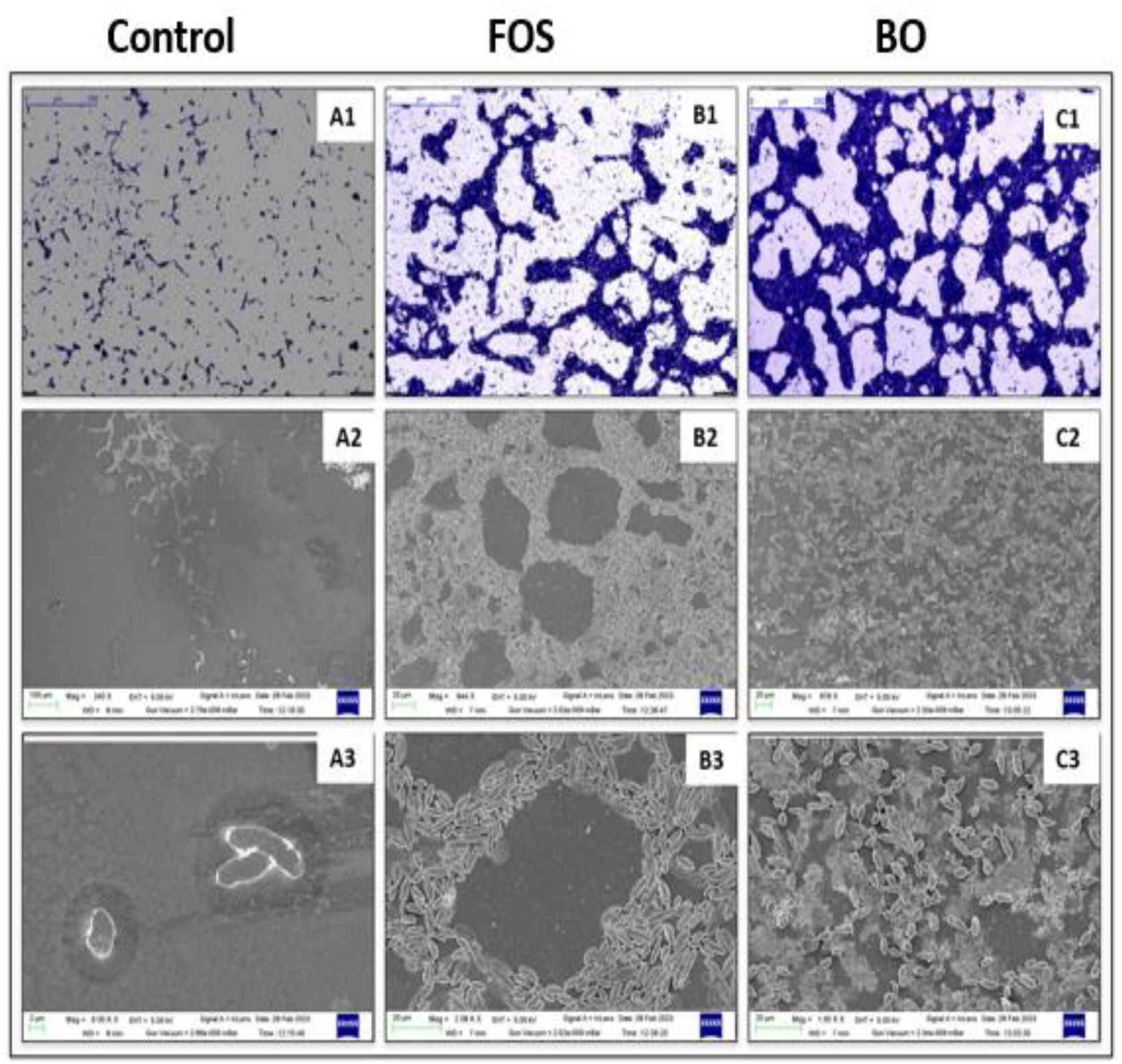
Representative images from the probiotic biofilm assay under three treatments: Control, FOS, and BO. Top row (A1–C1): bright-field microscopy images of biofilms formed by *Lactobacillus* spp. (scale bar = 250 µm). A1, Control (carbohydrate-free medium); B1, FOS; C1, BO. Middle row (A2–C2): SEM images illustrating surface attachment and biofilm architecture. Bottom row (A3–C3): higher-magnification SEM images highlighting bacterial clustering and biofilm organisation. Scale bars are indicated within individual panels.

SEM analysis corroborated these findings (Fig. 5A2–C3). Biofilms mediated by BO were dense and multilayered, characterised by a prominent extracellular matrix, reflecting enhanced cellular aggregation and surface adherence. FOS also promoted biofilm development; however, the resulting structures were thinner and less cohesive. Control samples displayed minimal adherence, with isolated cells and scattered microcolonies.

Quantitative measurements (Fig. 6) supported the microscopy findings. Biofilm biomass reached 391.33 ± 26.08% (relative to control) for BO, which was significantly higher than that observed for FOS (299.00 ± 19.29%) and the control (100%). The number of viable cells followed a similar trend: BO-supported biofilms contained 9.01 ± 0.70 log CFU/mL, whereas FOS and control contained 8.00 ± 0.69 and 3.88 ± 0.46 log CFU/mL, respectively. These results indicate that BO enhances biofilm biomass and cell retention under sessile growth conditions.

**Fig. 6.**
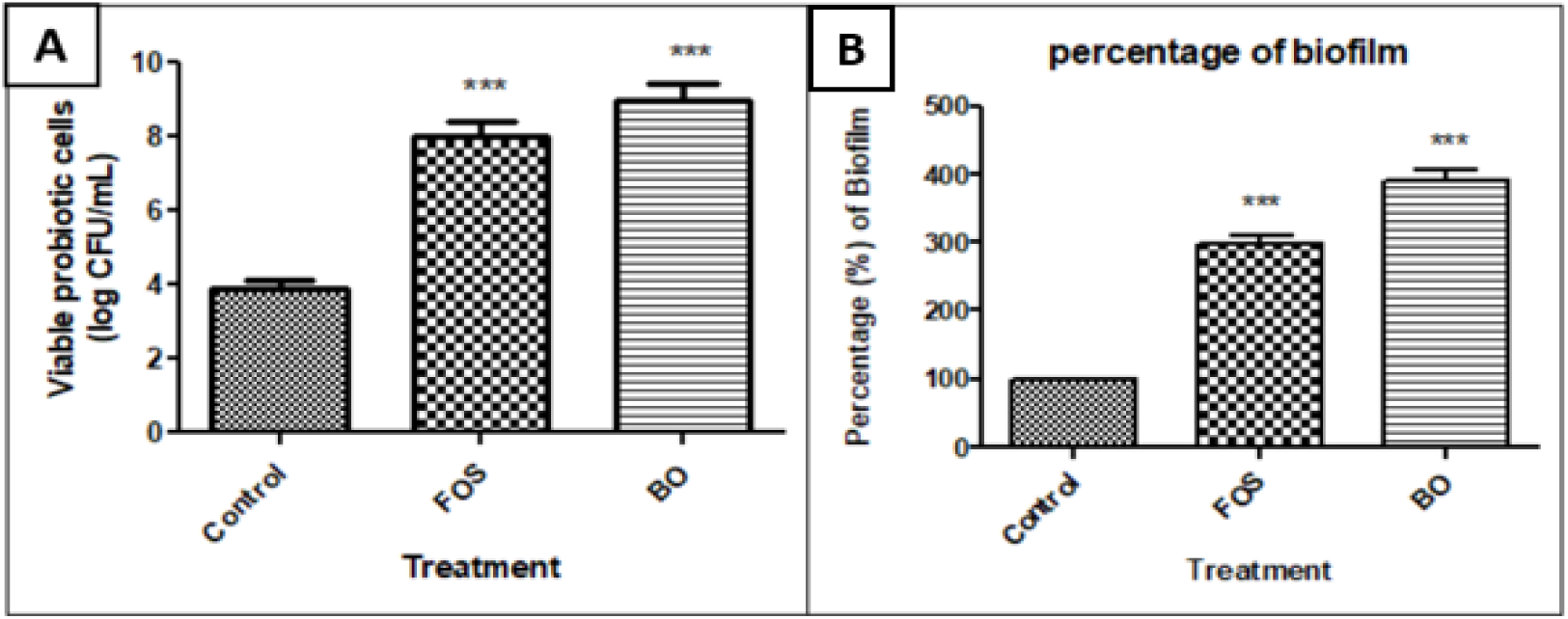
Quantitative analysis of probiotic biofilms: (A) viable cell counts (log CFU/mL) and (B) percentage biofilm formation. Results are expressed as mean values with corresponding SEM (n = 3). Differences among groups were analysed by one-way ANOVA followed by Dunnett’s multiple comparisons test (versus control). *p < 0.05, **p < 0.01, ***p < 0.001; # indicates not significant (p > 0.05) versus the control

The pronounced biofilm response observed with BO is consistent with reports showing that specific oligosaccharides can stimulate bacterial aggregation, extracellular matrix production, and surface colonisation in *Lactobacillus* species (Yin et al., 2024). Biofilm formation is a recognised determinant of probiotic functionality, as it can facilitate mucosal adherence and persistence, support epithelial barrier function, and contribute to pathogen exclusion (Jefferson, 2004; Lebeer et al., 2007). The use of a mixed probiotic culture comprising only probiotic strains enabled a focused assessment of BO-specific effects without interference from competitive microbial species, consistent with earlier colonisation studies (Jiang et al., 2023).

The lack of intricate three-dimensional formations, such as mushroom-shaped structures is not surprising, as these morphologies are predominantly associated with pathogenic biofilms developed under flow conditions and are seldom documented for Lactobacillus biofilms produced under static incubation (Jara et al., 2022; Martinez et al., 2020). The denser EPS layers and cohesive cellular aggregates found in BO-treated samples are well recognized as indications of mature probiotic biofilm development.

Collectively, these imaging and quantitative analyses indicate that BO was markedly more effective than FOS in facilitating substantial *Lactobacillus* spp. biofilm formation. The augmented structural integrity and enhanced viable biomass suggest superior probiotic adherence and sessile persistence. However, the results of the present study indicate that BO has the potential to support probiotic persistence and colonisation, which should be further evaluated in dynamic gut models to determine whether these biofilm-enhancing effects are maintained under physiologically relevant conditions.

## Conclusion

Black rice flour yielded an oligosaccharide-rich extract of 4.95 ± 0.33% (w/w) following extraction with 80% (v/v) aqueous ethanol at 60 °C for 1 h using a solvent-to-solute ratio of 5:1 (v/w). Subsequent purification through charcoal–celite chromatography followed by dialysis using a 500 Da MWCO membrane produced a structurally clean BO fraction of 2.87 ± 0.29% that was effectively devoid of mono-and disaccharides.

MALDI-TOF profiling and complementary chromatographic analyses demonstrated that the purified BO fraction consisted of a homologous series of hexose-based oligosaccharides with degrees of polymerisation ranging from 3 to 8 (m/z 527–1330). Acid hydrolysis followed by GC–MS identified glucose and galactose as detectable monomeric residues, while ketose-specific staining of the intact oligosaccharide fraction confirmed the presence of ketosyl residues, most plausibly attributed to fructosyl units based on cereal origin, DP distribution, and mass increment patterns, despite their disappearance following hydrolysis and derivatisation.

BO exhibited high digestive stability, with minimal hydrolysis under simulated salivary (2.08 ± 0.51%), gastric (0.34 ± 0.03%), and intestinal (4.29 ± 0.73%) conditions, leaving approximately 93% of the oligosaccharide fraction intact and available for colonic fermentation. The purified BO selectively stimulated probiotic *Lactobacillus* strains, showing the greatest prebiotic activity responses in *L. rhamnosus* (1.165 ± 0.255), *L. plantarum* (0.980 ± 0.163), and *L. casei* (0.944 ± 0.076). Fermentation of BO generated physiologically relevant SCFA with strain-dependent profiles.

In vitro biofilm assays, BO greatly promoted probiotic biofilm formation by enhancing the biofilm biomass to approximately 31% higher than that observed for FOS and supporting dense, matrix-rich biofilm architectures under static incubation. The increased sessile growth and structural integrity observed under BO supplementation suggest enhanced probiotic persistence and colonisation potential *in vitro*.

Overall, BO constitutes a digestion-resistant, structurally defined, and metabolically active oligosaccharide fraction of cereal origin that displays selective prebiotic utilisation, SCFA production, and substantial stimulation of probiotic biofilm formation. These findings establish BO as a promising prebiotic substrate and warrant further investigation using dynamic gut fermentation systems and *in vivo* models to evaluate functional efficacy under physiologically relevant gastrointestinal environments.

## Credit Authorship Contribution Statement

**Shivangi Agrawal:** Conceptualization, Methodology, Investigation, Formal analysis, Writing – original draft. **Paramita Biswas:** Investigation.**Susmita Mondal:** Investigation. **Abinaya Balasubramanian:** Investigation. **Sachin Maji:** Investigation. **Sandip Shit:** Investigation. **Satyabrata Ghosh:** Investigation. **Satyahari Dey:** Conceptualization, Project administration, Supervision, Writing – review & editing.

## Acknowledgements

Financial support from the Ministry of Education, Government of India, is gratefully acknowledged. We also acknowledge the Central Research Facility (CRF), IIT Kharagpur, for providing the necessary infrastructural and analytical support.

## Appendix

**ANOVA**: analysis of variance

**ATCC**: American Type Culture Collection

**BO**: black rice oligosaccharides

**CFU**: colony-forming units

**Da**: Dalton (unit)

**DHB**: 2,5-dihydroxybenzoic acid

**DNS**: 3,5-dinitrosalicylic acid (reagent)

**DP**: degree of polymerisation

**DP3–DP8**: degree of polymerisation 3–8

**EPS**: extracellular polymeric substance(s)

**FOS**: fructo-oligosaccharides

**GC–MS**: gas chromatography–mass spectrometry

**GIT**: gastrointestinal tract

**GOS**: galacto-oligosaccharides

**HPLC**: high-performance liquid chromatography

**i.d.**: internal diameter

**M9**: M9 minimal medium

**min**: minute(s)

**m/z**: mass-to-charge ratio

**MALDI-TOF MS**: matrix-assisted laser desorption/ionization time-of-flight mass spectrometry

**MRS**: de Man, Rogosa and Sharpe (medium)

**MSTFA**: N-methyl-N-(trimethylsilyl)trifluoroacetamide

**MTCC**: Microbial Type Culture Collection

**MWCO**: molecular weight cut-off

**NCDC**: National Collection of Dairy Cultures

**NIST**: National Institute of Standards and Technology

**PBS**: phosphate-buffered saline

**RT**: retention time

**SCFAs**: short-chain fatty acids

**SEM**: scanning electron microscopy

**spp.**: multiple species (plural of sp.)

**TLC**: thin-layer chromatography

